# Genome-wide chemical mutagenesis screens allow unbiased saturation of the cancer genome and identification of drug resistance mutations

**DOI:** 10.1101/066555

**Authors:** Jonathan S. Brammeld, Mia Petljak, Inigo Martincorena, Steven P. Williams, Luz Garcia Alonso, Alba Dalmases, Beatriz Bellosillo, Carla Daniela Robles-Espinoza, Stacey Price, Syd Barthorpe, Patrick Tarpey, Constantine Alifrangis, Graham Bignell, Joana Vidal, Jamie Young, Lucy Stebbings, Kathryn Beal, Michael R Stratton, Julio Saez-Rodriguez, Mathew Garnett, Clara Montagut, Francesco Iorio, Ultan McDermott

## Abstract

Drug resistance is an almost inevitable consequence of cancer therapy and ultimately proves fatal for the majority of patients. In many cases this is the consequence of specific gene mutations that have the potential to be targeted to re-sensitize the tumor. The ability to uniformly saturate the genome with point mutations without chromosome or nucleotide sequence context bias would open the door to identify all putative drug resistance mutations in cancer models. Here we describe such a method for elucidating drug resistance mechanisms using genome-wide chemical mutagenesis allied to next-generation sequencing. We show that chemically mutagenizing the genome of cancer cells dramatically increases the number of drug-resistant clones and allows the detection of both known and novel drug resistance mutations. We have developed an efficient computational process that allows for the rapid identification of involved pathways and druggable targets. Such *a priori* knowledge would greatly empower serial monitoring strategies for drug resistance in the clinic as well as the development of trials for drug resistant patients.

## Introduction

Despite an increasing array of new cancer therapies, drug resistance is an almost universal phenomenon that is likely due to the presence of rare subclonal populations that act as a reservoir for resistance mutations. The emergence of drug resistance ultimately proves fatal for the majority of patients, and therefore the early detection of resistance and the identification of novel re-sensitization strategies is a subject of intense activity.

Previously, the identification of drug resistance genes has relied on either re-biopsy of cancer patients following the development of resistance or the use of cancer cell lines made resistant by exposure *in vitro* to drug over many weeks. Both approaches can suffer from inherent biases. With respect to the former, biopsy of a single resistant site of disease may miss alternate resistance mechanisms in other metastatic sites (Van Allen et al. 2014). Equally, serial drug exposure in cancer cell lines will favor pre-existing drug resistant clones that are specific for that cell line and may not represent the entire spectrum of resistance mechanisms for that treatment.

For these reasons, there is considerable interest in the use of forward genetic screens capable of engineering into the cancer genome mutational events that can be tested for their ability to cause drug resistance in an unbiased fashion. Such screens, if sufficiently unbiased, could in theory capture the entire breadth of genetic resistance mechanisms for any drug. Recent studies have demonstrated the power of both genome-wide gain- and loss-of-function screens using CRISPR/Cas9, lentiviral shRNA and large-scale open-reading frame technologies to identify clinically relevant drug resistance mechanisms in cancer (Hu and Zhang 2016). However, these screens all fail to capture a third important mechanism of drug resistance, namely that of point mutations. Point mutations account for resistance in large numbers of patients receiving targeted therapies in melanoma, colon and lung cancers and chronic myeloid leukemia (**Supplemental Table S1**) (Kobayashi et al. 2005; Katayama et al. 2012; Montagut et al. 2012; Ohashi et al. 2012; Bettegowda 2014; Long et al. 2014; Van Allen et al. 2014; Wagle et al. 2014; Arena et al. 2015; Russo et al. 2015; Siravegna et al. 2015; Thress et al. 2015).

N-Ethyl-N-nitrosourea (ENU) has been used as a potent mutagen in mouse models of development for over four decades (Acevedo-Arozena et al. 2008). Exposure results in the efficient generation of random point mutations throughout the cell genome (Tokunaga et al. 2014). We therefore tested whether, in cancer cell line models, ENU could be used to mutagenize the genome and enable expansion of drug resistant cells following the application of a targeted agent. As proof of concept, we chose to investigate whether this approach could identify all clinically demonstrated resistance mutations in colorectal cancer patients treated with the EGFR monoclonal antibody Cetuximab (Van Cutsem et al. 2009). In the clinic, resistance in such patients is heavily driven by point mutations and a decade of clinical studies has identified the vast majority of the resistance mutations. This ‘ground truth’ should in theory allow us to define how well a saturation mutagenesis screen can identify clinically relevant resistance mutations.

We developed a sequencing and informatics approach to detect novel resistance mutations from next-generation sequence data and to detect statistical enrichment for mutually exclusive mutations in specific signalling pathways comprising more than 8,000 genes at the sample population level. Our mutagenesis screen was able to successfully identify all known drug resistance mutations to Cetuximab previously observed in the clinic as well as a novel mutation that we subsequently identified in a colorectal cancer patient. We suggest that this approach is a powerful and facile means to draw the landscape of point mutations that confer resistance to targeted therapies. Such knowledge could be used to discover therapeutic strategies to re-sensitize resistant tumors as well as identify which genes should be prioritised for non-invasive monitoring during treatment using plasma DNA sequencing.

## Results

### ENU exposure confers stable resistance to Cetuximab in colon cancer cells

We screened 51 colorectal cancer cell lines with a concentration range of the EGFR monoclonal antibody Cetuximab and assessed viability after 6 days (Figure 1A). In keeping with clinical experience of the genetic factors that underpin response to this drug, those cell lines wild-type for *KRAS*/*NRAS*/*BRAF* (green bars) exhibited heightened sensitivity to Cetuximab (Douillard et al. 2013). We therefore chose two of these lines, CCK-81 and NCI-H508, to use in the ENU resistance experiment. Both cell lines additionally demonstrated Cetuximab sensitivity in long-term clonogenic survival assays (**Supplemental Fig. S1**). Moreover, CCK-81 has features of microsatellite instability (MSI) whereas NCI-H508 is microsatellite stable (MSS). MSI is detected in 16% of colorectal cancers and is associated with a different phenotype and clinical outcome compared to MSS cancers. The CCK-81 cell line was exposed to a dose range of ENU (0.01-1mg/ml) for 24 hours, following which the mutagenized cells were treated with Cetuximab (10µg/ml) for 8 consecutive weeks. The number of drug resistant colonies was counted at the end of the experiment. Importantly, we observed no drug resistant colonies in the absence of ENU (Figure 1B). With increasing ENU concentration we observed a linear increase in both the number of drug resistant colonies (left y-axis, blue bars) as well as the number of mutations per clone (right y axis, green triangles). We subsequently used a concentration of ENU (0.1mg/ml) that resulted in minimal viability effect in both cell lines (see Methods). We next treated NCI-H508 cells with ENU (0.1mg/ml) for 24 hours followed by weekly Cetuximab treatment for 8 weeks. Drug resistant colonies were picked, expanded in culture and 72 were submitted for whole exome Illumina sequencing (a total of 14 CCK-81 and 58 NCI-H508 colonies). Data was analysed for substitutions and insertions/deletions to enable an estimation of the number of ENU-associated mutations per Mb of exome and to detect novel (and putative drug resistance) mutations (**Supplemental Table S2**). We then performed clonogenic survival assays on a subset of resistant clones and confirmed robust and stable resistance to Cetuximab (Figure 1C).

**Figure 1.**
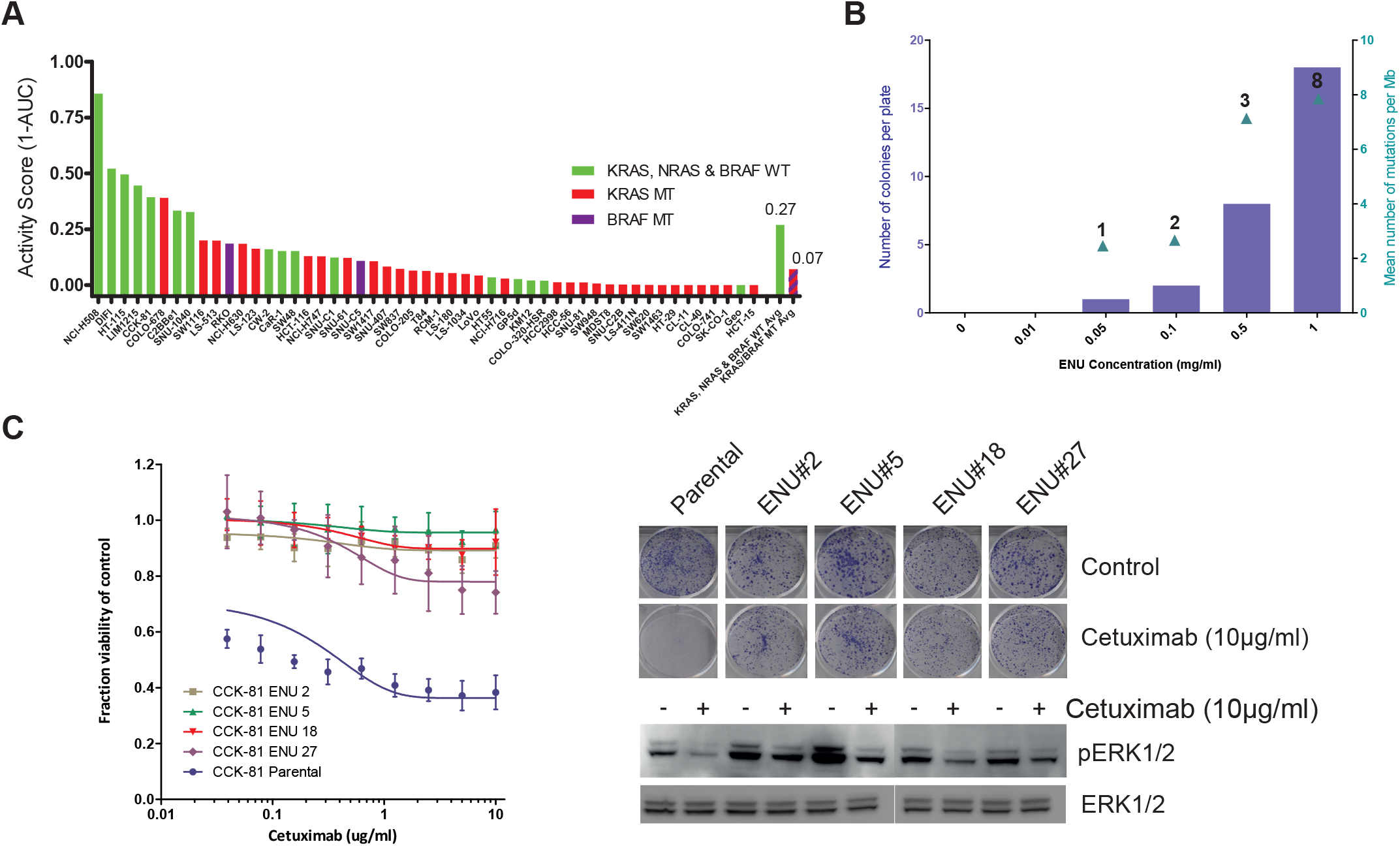
Genome-wide chemical mutagenesis screens to define pathways of drug resistance in cancer. (A) Cetuximab viability screen in colorectal cancer cell lines. 51 colorectal cancer cell lines were screened with a concentration range of the EGFR monoclonal antibody Cetuximab and viability (as measured by the Activity Score, 1-AUC) was measured after 6 days. The KRAS, NRAS and BRAF mutation status of each line is indicated. The mean Activity Score for those cell lines wild-type for all 3 genes (green bar) versus those with a canonical driver mutation in at least one gene (red bar) is indicated in the last two columns. (B) Dose dependent effect of ENU on mutation burden and drug resistance. CCK-81 colorectal cancer cells were treated for 24 hours with increasing concentrations of ENU (x-axis, 0.01-1mg/ml) and then weekly with Cetuximab (10µg/ml) for 8 weeks to allow drug resistant colonies to develop. The number of resistant colonies per plate were counted for each ENU concentration (blue bars) and submitted for whole exome sequencing to calculate the mean number of mutations per Mb (green triangles). Numbers above each triangle indicate the number of clones sequenced at that concentration. (C) ENU mutagenesis generates stably drug-resistant clones. A subset of Cetuximab-resistant clones generated in the CCK-81 cell line following exposure to ENU and subsequent serial weekly Cetuximab treatment were picked from the plate and expanded separately out of drug for 4 weeks. 21-day clonogenic survival assays in 4 clones (along with the parental CCK-81 cell line) treated with Cetuximab 10µg/ml confirmed that resistance to the EGFR mAb had been maintained and was stable. A 6-day viability assay of cells treated with a concentration range of Cetuximab (right panel) demonstrates resistance of ENU clones at all concentrations. Immunoblot analysis of effect of Cetuximab treatment (10µg/ml) for 6 hours confirms persistence of MAPK signalling in ENU clones.

### The spectrum of ENU-induced mutations

The ability of any mutagenesis screen to capture a particular phenotype is strongly dependent on its ability to evenly saturate the genome with all 6 possible classes of base substitution type (expressed as the pyrimidine of a mutated Watson–Crick base pair, C>A, C>G, C>T, T>A, T>C, T>G). On average, we detected 470 novel mutations per exome in each clone (mean 570 and 446 in CCK-81 and NCI-H508 clones respectively), for a total of 33,857 (**Supplemental Table S2**). The mutations were almost exclusively composed of base substitutions (96% of total). A third of such mutations were non-synonymous (missense) variants within the coding exon of a gene where resistance mutations are more likely to occur (Figure 2A). Only 4% were potential loss-of-function truncating mutations (frameshift indels or nonsense mutations). The remaining mutations were predominantly either silent or intronic. Significantly, analysis of exome sequence data across all 72 clones (regardless of whether MSS or MSI) revealed that of the 6 possible classes of base substitution, only C>G substitutions are less well represented (3% of all substitution base changes) (Figure 2A). The mutation spectrum in ENU-derived clones was similar regardless of whether the cells came from a MSI or MSS background (**Supplemental Fig. S2**). There was no evidence of a significant bias towards mutations in coding genes in any particular chromosome or indeed any specific region within a chromosome (Figure 2B **and Supplemental Fig. S3**). However, given these 33,857 mutations are an admixture of those caused by ENU, background mutational processes and private subclonal variants, we elected to use a mathematical approach to specifically extract the ENU signature from the data and more accurately determine the mutation spectrum of ENU mutations.

**Figure 2.**
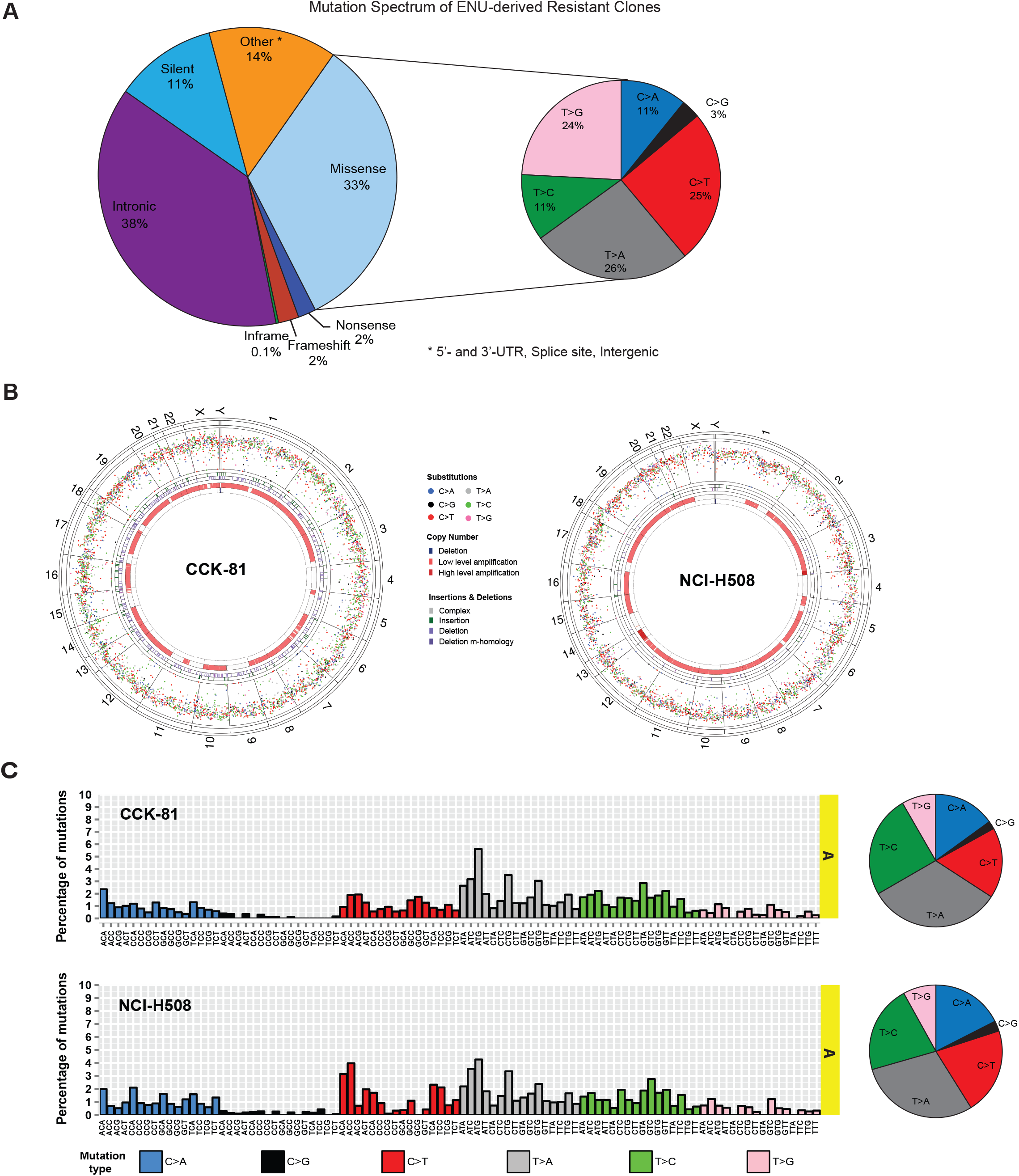
The spectrum of ENU-induced mutations. (A) Spectrum of mutations in ENU-derived drug resistant clones. A pie chart representation of the proportion of 33,857 mutations detected in CCK-81 and NCI-H508 clones categorised according to mutation type. For the missense mutations, there is a further representation of the proportion of mutations falling within the 6 possible nucleotide base substitutions (C>A, C>G, C>T, T>A, T>C, T>G). (B) Circos plots of mutations in CCK-81 and NCI-H508 clones. All substitution mutations in CCK-81 or NCI-H508 clones are represented by their intermutation distance per chromosome. Each chromosome is composed of all coding exons in genes found within that chromosome. The four circles in each plot, from outermost inwards, represent the human chromosomes, substitution mutations, insertion mutations, deletion mutations, copy number gains, and copy number losses. Low and high level amplification refers to 2-4 and 4-8 copies respectively. (C) The mutational signature of ENU. The trinucleotide representation of ENU mutations in CCK-81 and NCI-H508 drug-resistant colonies is displayed as the distribution of mutations for all 96 possible combinations of mutations generated as a result of ENU exposure. Adjacent pie charts display the relative distribution of each of the six classes of possible base substitutions for the CCK-81 and NCI-H508 clones.

The non-negative matrix factorization algorithm has previously been used to detect the presence of mutational signatures in human cancers, including from defects in DNA mismatch repair and altered activity of the error-prone polymerase *POLE* (http://cancer.sanger.ac.uk/cosmic/signatures) (Alexandrov et al. 2013b). It extracts signatures based on a 96-mutation classification that incorporates the 6 base substitution types described above as well as the immediate flanking sequence context of the mutated base (four possible 5’ and four possible 3’ bases). In our data it revealed a distinct and unique signature that was represented across almost all trinucleotide contexts in both CCK-81 and NCI-H508 clones and not previously detected in prior tumor studies, including a panel of 51 colorectal cancer cell lines (data not shown) (Figure 2C, **Supplemental Fig. S4**). This signature (‘Signature A’) is likely one of ENU exposure. Reassuringly, the pattern of base substitutions that comprise this signature was almost identical to that seen across the entire set of substitutions detected in the ENU-derived clones, with again only C>G substitutions seen at lower frequency (Figure 2C). Thus, using this approach it should be feasible to generate the majority of theoretical coding point mutations for drug resistance across the entire genome.

As expected we detected a signature of MSI (‘Signature B’) in the combined mutational catalogue for the CCK-81 clones but surprisingly also in the NCI-H508 clones (**Supplemental Fig. S4**) (Alexandrov et al. 2013a). On closer examination, this signature was the result of two hypermutator clones in the NCI-H508 mutational catalogue (red arrows) (clones NCI-H508_26 and NCI-H508_40) (**Supplemental Fig. S5**). These clones had mutation rates as high as any of the MSI CCK-81 clones and increased numbers of small insertions and deletions. This would be in keeping with a defect in the mismatch repair pathway (**Supplemental Table S2**). Clone NCI-H508_26 was found to harbor a novel nonsense (stop-gained) mutation in the mismatch repair gene *MLH1* and clone NCI-H508_40 a nonsense mutation in the DNA repair gene *EXO1*. Two other clones (black arrows) also have elevated mutation rates that may be the result of gaining nonsense mutations in *POLQ*, a gene involved in DNA damage repair. These gave rise to the third signature of unknown origin detected in NCI-H508 clones (‘Signature C’) (**Supplemental Fig.S4**).

### ENU mutagenesis identifies clinically relevant resistance mutations and pathways

A challenge in the identification of drug resistance mutations in ENU-derived clones is that each clone harbors many hundreds of ‘passenger’ mutations in addition to that conferring resistance. We hypothesised that with a sufficient population of individual resistant clones it might become feasible to use statistical enrichment for non-synonymous coding mutations in specific pathways to help identify drug resistance mutations. We therefore employed a statistical framework (SLAP-Enrich) to identify whether genetic alterations observed in multiple samples are enriched within a specific pathway in a statistically significant manner using a network of 8,056 unique genes (https://github.com/saezlab/SLAPenrich) (lorio et al. 2016). Once significantly enriched pathways are identified SLAP-Enrich applies a final filter based on the tendency for genes in a positively selected pathway to be mutated in a mutually exclusive manner. When applying this method to the set of ENU-mutations across the 72 Cetuximab-resistant clones, we found several statistically enriched pathways (false discovery rate (FDR) < 5%) (**Supplemental Table S3**). The pathway most significantly enriched with mutations, ‘Signalling to P38 via RIT and RIN’, contains many of the key genes of the canonical MAP kinase pathway (Figure 3A). In total, we were able to identify credible resistance mutations in 42 of the 72 resistant clones (59%) (**Supplemental Table S4**). We detected credible resistance mutations in all of the genes previously found clinically to confer resistance to EGFR therapy in colorectal cancer (**Supplemental Table S1**). *EGFR*, *KRAS*, *NRAS*, *BRAF* and *MAP2K1* (*MEK1*) were each found to be mutated in ≥3 clones and in a mutually exclusive manner. There was no clear difference in the frequency of specific mutations between NCI-H508 and CCK-81. - Furthermore, 38/42 (90%) of these putative resistance mutations have previously been identified in colorectal patients developing resistance to Cetuximab. The most frequently observed ENU resistance mutation was that of *BRAF* p.V600E (13/42 clones), followed by *NRAS* p.Q61K (8/42) and *KRAS* p.G12C (4/42) (Figure 3B,C and **Supplemental Fig. S6**). These mutations are all canonical driver mutations in tumorigenesis and known to activate oncogenic signalling and confer resistance to Cetuximab both experimentally and clinically (Diaz et al. 2012; Misale et al. 2012). We also detected *EGFR* mutations in 3 of the resistant clones. The *EGFR* I491K substitution has been shown to induce structural changes to the extracellular domain of EGFR such as to prevent Cetuximab binding and confer resistance (Montagut et al. 2012). Of note, additional mutations also present in the ‘Signalling to P38 via RIT and RIN’ network (e.g. JAK1, IL6R, RAF1) were not taken forward for investigation as these mutations were present in clones also harbouring other more credible drivers of drug resistance and were present at low frequency (<3 clones) (**Supplemental Fig. S7**).

**Figure 3.**
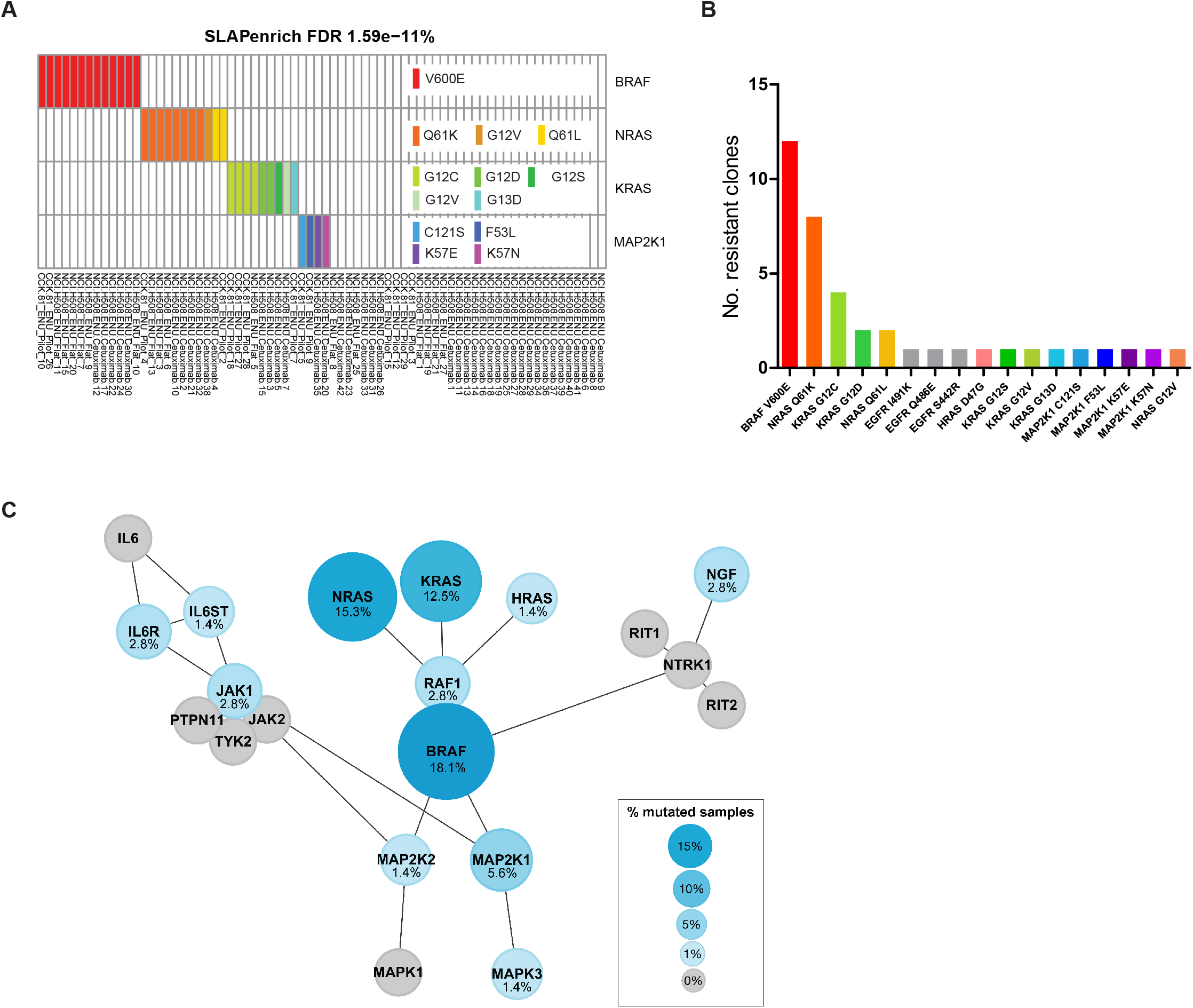
ENU mutagenesis identifies clinically relevant mutations and pathways. (A) The pathway most significantly enriched with mutations, ‘Signalling to P38 via RIT and RIN’, contains many of the key genes of the canonical MAP kinase pathway and which are mutated in a mutually exclusive manner. Here we show those genes mutated in at least 3 individual clones. Whole exome sequence from 72 Cetuximab resistant ENU clones was used to identify pathways using the SLAP-Enrich algorithm. Amino acid substitutions are labelled for the subset of genes most frequently mutated in the pathway and/or demonstrating mutual exclusivity with other mutations. (B) Cetuximab resistance mutations. Frequency of likely ENU-derived drug resistance mutations across 42 resistant clones. The amino acid consequence of each mutation is indicated with the gene name. (C) A visual representation of the all mutated genes that comprise the ‘Signalling to P38 via RIT and RIN’ pathway indicating gene-gene interactions and the hierarchy of signalling. Only those genes mutated in at least 3 clones were taken forward for validation.

### A mutation enrichment analysis identifies drug resistance genes

Recent studies of large mutational datasets from cancer sequencing studies have used statistical approaches that consider the mutation spectrum, the sequence of each gene, the impact of coding substitutions (synonymous, missense, nonsense, splice site) and the variation of the mutation rate to detect novel cancer genes (Nik-Zainal et al. 2016). We adapted this ‘*d*_N_/*d*_S_’ (nonsynonymous/synonymous) method to analyse the mutations identified in the ENU drug resistant clones (see Methods). As this approach was designed for the analysis of unrelated tumor samples and we have instances of clones sharing mutations that reflect a common subclonal origin within the original parental cell lines, we first condensed the 72 ENU clones into 19 representative groups, each the union of all the mutations in clones sharing more than 3 mutations. Three genes were identified as having a pattern of mutations supporting being under positive selection, namely *NRAS*, *KRAS* and *MAP2K1* (FDR < 0.05; **Supplemental Table S5**). After these the next highest ranked gene was *BRAF*; although it was of borderline significance (FDR=0.0508), it is a very strong candidate for a resistance gene given that 4/6 of the mutations are at the canonical p.V600 locus.

### ENU mutagenesis identifies novel resistance mutations in MAP2K1

Recently, two studies of plasma DNA sequencing in colorectal cancer patients undergoing treatment with EGFR monoclonal antibodies jointly identified the first *MAP2K1* (*MEK1*) codon K57 resistance mutations (p.K57T and p.K57N) (Russo et al. 2015; Siravegna et al. 2015). In our study we also identified *MAP2K1* mutations at the K57 codon (p.K57N, p.K57E) as well as at two sites not previously reported (p.F53L, p.C121S) (**Supplemental Table S4**). We therefore sequenced these *MAP2K1* loci (together with additional mutation hotspots in 34 other genes) in a series of plasma DNA samples collected from 22 colorectal cancer patients who acquired resistance to treatment with EGFR therapies (either Cetuximab or Panitumumab) after an initial response. In addition to all of the known canonical resistance mutations (in *KRAS*, *NRAS* and *BRAF*), we detected in one such patient a novel p.F53L *MAP2K1* mutation predicted by our screen to be a resistance mutation (**Table 1**, **Figure 4A**). As previously reported, we detected in a number of these patients more than one likely resistance mutation, in keeping with different metastatic sites evolving different resistance mechanisms.

**Table 1.** Identification of genetic alterations associated with resistance to anti-EGFR antibodies in plasma samples from 22 colorectal cancer patients. The table lists putative genetic mechanisms of acquired resistance to anti-EGFR therapies that were identified in ctDNA of 22 patients. The novel MAP2K1 mutation is indicated in red. FOLFIRI = folinic acid, 5-fluorouracil and irinotecan; FOLFOX = folinic acid, 5-fluorouracil and oxaliplatin. ND = no resistance mutation detected.

| Patient | Therapy | Resistance | Resistance mutation | Mutant Allele (%) | Oncogenic alteration in COSMIC | Druggable |
| --- | --- | --- | --- | --- | --- | --- |
| 1 | FOLFIRI + Cetuximab | Acquired | KRAS p.Q61H | 47% | YES | NO |
| 2 | FOLFIRI + Cetuximab | Acquired | ERBB2 amplification | - | YES | YES |
| 3 | FOLFIRI + Cetuximab | Acquired | KRAS p.G12V<br>MAP2K1 p.F53L | 36%<br>14% | YES<br>YES | NO<br>YES |
| 4 | Irinotecan+Cetuximab | Acquired | ND | - | - | - |
| 5 | Irinotecan+Cetuximab | Acquired | KRAS p.Q61H | 61% | YES | NO |
| 6 | Irinotecan+Cetuximab | Acquired | ND | - | - | - |
| 7 | FOLFOX + Cetuximab | Acquired | EGFR p.S492R | 1% | - | YES |
| 8 | FOLFOX + Cetuximab | Acquired | KRAS p.G12S | 14% | YES | NO |
| 9 | Irinotecan + Cetuximab | Acquired | KRAS p.G12A<br>NRAS p.Q61H | 6%<br>11% | YES<br>YES | NO<br>NO |
| 10 | Irinotecan + Cetuximab | Acquired | KRAS p.A146T | 26% | YES | NO |
| 11 | FOLFOX+ Cetuximab | Acquired | NRAS p.Q61L<br>EGFR p.S492R | 31%<br>6% | YES<br>- | NO<br>YES |
| 12 | Irinotecan + Cetuximab | Acquired | ND | - | - | - |
| 13 | Irinotecan + Cetuximab | Acquired | BRAF p.V600E | 14% | YES | YES |
| 14 | Irinotecan + Cetuximab | Acquired | NRAS p.Q61K | 35% | YES | NO |
| 15 | Irinotecan + Cetuximab | Acquired | EGFR p.R451C | 6% | - | - |
| 16 | Irinotecan + Cetuximab | Acquired | ND | 13% | YES | YES |
| 17 | Panitumumab | Acquired | ND | - | - | - |
| 18 | Irinotecan + Cetuximab | Acquired | EGFR p.S464L | - | - | YES |
| 19 | FOLFIRI + Cetuximab | Acquired | KRAS amplification | - | YES | - |
| 20 | FOLFIRI + Cetuximab | Acquired | ND | - | - | - |
| 21 | Cetuximab | Acquired | BRAF p.V600E | 6% | YES | YES |
| 22 | Irinotecan+Cetuximab | Acquired | EGFR p.K467T | 15% | - | - |

**Figure 4.**
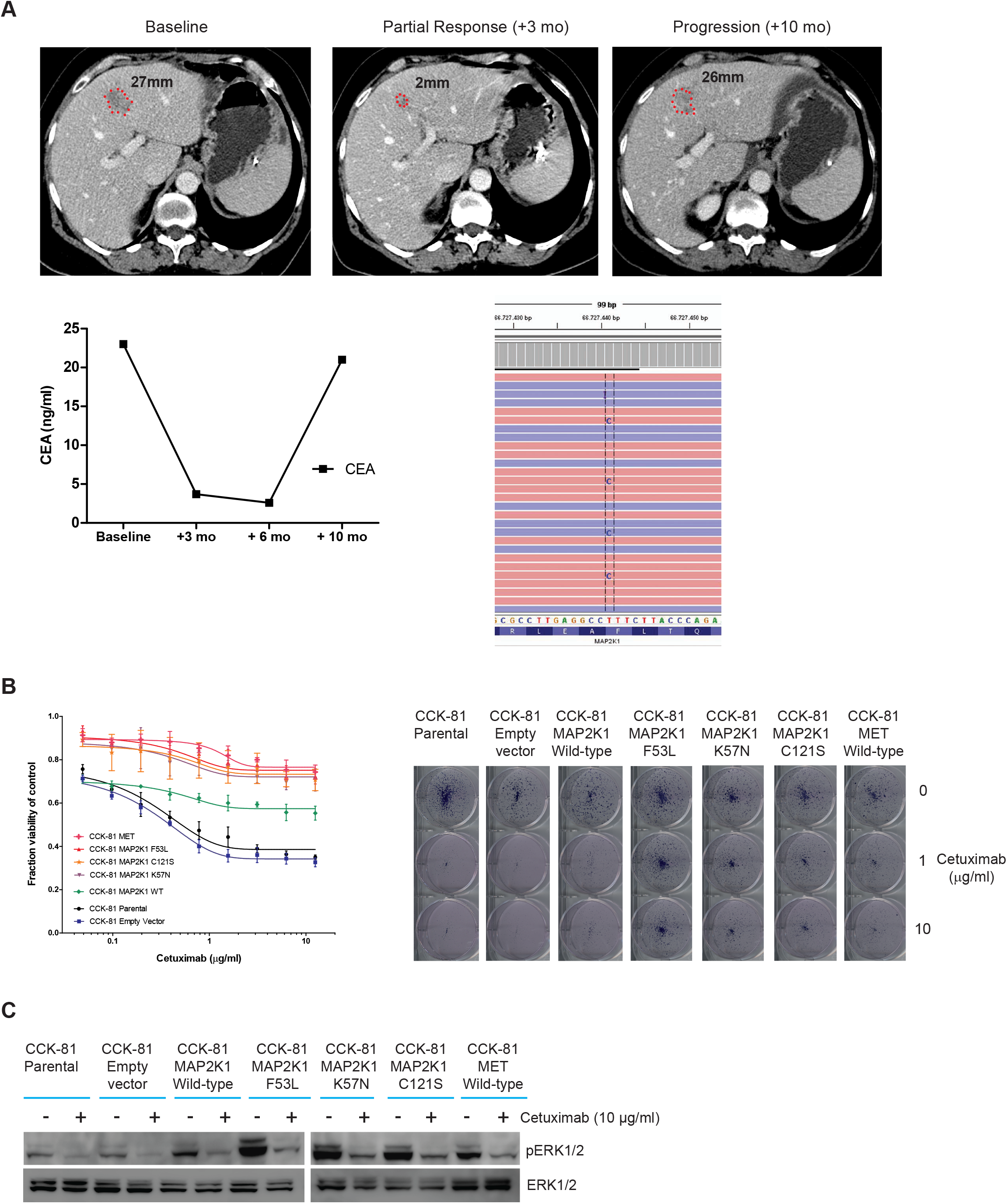
ENU mutagenesis identifies a novel MAP2K1 resistance mutation that is detected in a colorectal cancer patient after an initial response to Cetuximab. (A) A colorectal cancer patient with inoperable liver metastases was treated with the combination of FOLFIRI chemotherapy and Cetuximab. Initial response to treatment was demonstrated radiologically at 3 months and by falling CEA marker levels. Subsequent measurements at 10 months confirmed progressive disease and sequencing of plasma DNA detected a novel mutation in MAP2K1 (MEK1), p.F53L. (B) CCK-81 cells were transduced with the following lentiviral MAP2K1 vectors: empty vector (mock), wild-type (WT), K57N, C121S and F53L mutations. MET was also overexpressed as a positive control. Cells were seeded in 96-well plates and treated for 1 week with increasing concentrations of Cetuximab (0-12µg/ml) (left panel) or assayed using a clonogenic survival assay at 20 days at 1 or 10 µg/ml of Cetuximab (right panel). (C) Immunoblot of the effect of Cetuximab (10 µg/ml) at 6 hours on pERK in MAP2K1 mutant CCK-81 cells (as b).

To functionally validate the resistance effects of these *MAP2K1* mutations, we treated CCK-81 cells expressing the novel p.F53L and p.C121S mutations as well as the previously identified p.K57N mutation with Cetuximab (alongside empty vector and wild-type *MAP2K1* controls). We found that all of our candidate resistance mutations induced resistance to Cetuximab and the strength of the resistance effect for the mutations was comparable to that conferred by overexpression of the MET receptor tyrosine kinase, a previously identified resistance mechanism (Figure 4B, left panel) (Bardelli et al. 2013). Long-term growth inhibition assays similarly showed robust and durable resistance to Cetuximab in the *MAP2K1* mutant cells (Figure 4B, right panel). Immunoblot analysis demonstrated elevated constitutive phosphorylation of ERK1/2 as well as a failure to completely suppress pERK1/2 expression following Cetuximab treatment in all of the MAP2K1 clones (Figure 4C).

### Rational targeting of pathways can re-sensitize drug resistant mutants to Cetuximab

Constitutive EGFR signalling in solid tumors activates a number of downstream pro-survival/proliferation pathways including PKC, PI3K/AKT/mTOR, JAK-STAT and MAPK (**Supplemental Fig. S8A**) (Shostak and Chariot 2015). In EGFR-dependent cells, treatment with EGFR inhibitors affects cell survival by shutting down such processes. Identification of the key signalling pathways that underpin drug resistance opens up the possibility of rationally targeting key components of such resistance pathways and thus re-sensitizing cells. The creation of mutagenized resistant cell lines, either through the ENU screen or through the deliberate genetic modification of the parental cell line for specific mutations, allowed us the opportunity for such experiments to be carried out *in vitro*. As in clinical practice, the pathway most frequently mutated in the drug resistant CCK-81 and NCI-H508 ENU clones converges towards MAPK family members and targeting these nodes might be expected to overcome resistance (Figure 3C). For example, a Cetuximab-resistant CCK-81 *BRAF* V600E mutant clone (ENU-10) was re-sensitized when the EGFR monoclonal antibody was combined with the BRAF inhibitor Dabrafenib (**Supplemental Fig. S9A**). In the mutant cells the activating *BRAF* mutation would enable such cells to continue to signal through the MAPK pathway (and survive) despite EGFR blockade (**Supplemental Fig. S8B**). Thus, only by targeting both EGFR and BRAF in combination can all the relevant survival effectors be silenced and the viability of the cells reduced. Cetuximab-resistant clones harbouring mutations (in *KRAS, NRAS* and *MAP2K1*) that would be predicted to activate MAPK signalling were re-sensitized when a MEK inhibitor (Trametinib) was combined with Cetuximab (**Supplemental Fig. S9B**). Similarly, combining Cetuximab with Trametinib almost completely re-sensitized the resistant *MAP2K1* mutant CCK-81 cells (Figure 5A,B). Indeed, such a combination has already been suggested as putative therapeutic strategy for colon cancer patients (Misale et al. 2014).

**Figure 5.**
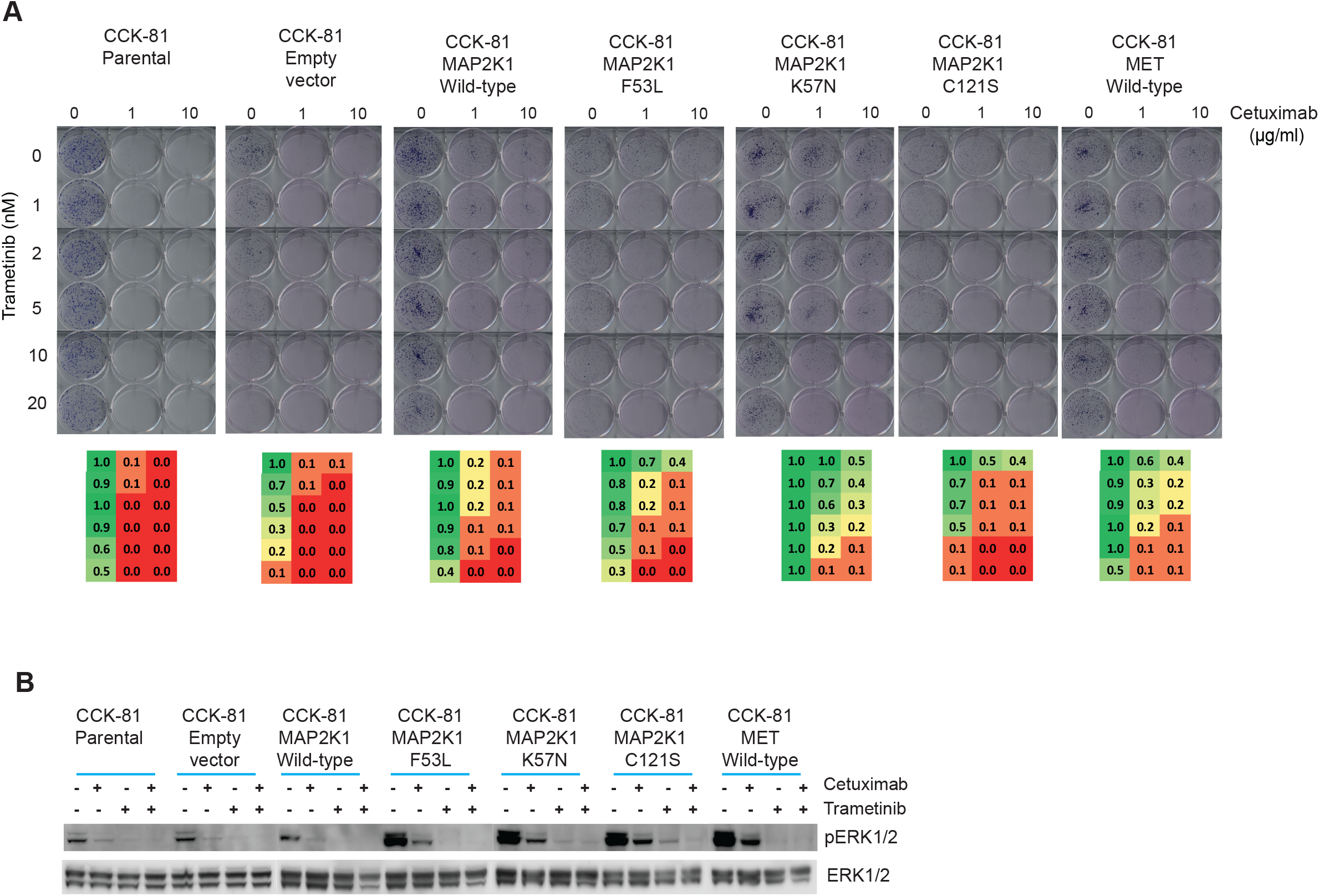
Rational targeting of resistance pathways to re-sensitize drug resistant mutants. (A) Clonogenic survival assay of MAK2K1 mutant CCK-81 cells when treated with Cetuximab, Trametinib or a combination of the two drugs. Heat maps of normalised viability effects per well are shown below each cell line. (B) Immunoblot of the effect of combing Cetuximab and Trametinib (MEK inhibitor) on MAPK signalling in MAP2K1 mutant CCK-81 cells. Cells were treated with Cetuximab (10 µg/ml), Trametinib (5nM) or both for 6 hours. MET was also overexpressed as a positive control.

## Discussion

We proposed at the onset of this study that regardless of what sequencing technology is used to detect resistance point mutations, some *a priori* knowledge of the likely drug resistance candidates should greatly increase the sensitivity of such assays. Identifying the complete catalogue of drug resistance effectors to any drug requires *in vitro* studies that model resistance in the relevant tissue and genetic background. In the past such *in vitro* studies featured cell lines that had undergone serial passage in the presence of the candidate drug in order to force the emergence of resistant clones (Ogino et al. 2007; Turke et al. 2010; Katayama et al. 2011; Eberlein et al. 2015). Although such studies have successfully identified clinically relevant mechanisms of drug resistance in some instances, they are biased towards selecting for those pre-existing resistant subclones that are particular to that specific cell line.

As a means of generating random point mutations throughout the genome, ENU chemical mutagenesis and subsequent phenotype-driven screening has been pivotal to a complete understanding of how complex biological processes operate in classical model organisms including yeast (Forsburg 2001), flies (St Johnston 2002), zebrafish (Patton and Zon 2001), and, perhaps most extensively, mice (Kile and Hilton 2005). The alkylating agent *N*-ethyl-*N*-nitrosourea (ENU) can introduce a high rate of point mutations into the genome and has two distinct advantages over previously used mutagens. First, it is very efficient, inducing a point mutation every 1 to 2 Mb throughout the genome in mouse models (~100-fold higher than the spontaneous mutation rate and 3-fold higher than X-irradiation) (Concepcion et al. 2004). Second, unlike irradiation, which induces multi-locus deletions, ENU is a point mutagen and affects single loci. ENU functions by the transfer of its ethyl group to Oxygen or Nitrogen atoms in DNA, resulting in misidentification of these ethylated bases during replication. If the mismatch is not repaired a base-pair substitution results (Justice 2000). To date, the use of ENU to define drug resistance mechanisms in cancer has been focused on specific genes in non-cancer cell line models rather than to interrogate the entire coding genome (Tiedt et al. 2011; Zhang et al. 2011; Ercan et al. 2015). Previous sequencing studies of ENU-derived mutations in mouse and fly models had demonstrated a strong bias in the spectrum of substitution mutations observed, with especially low numbers of C>G, T>G and C>A mutations. We observed a much more balanced representation of all 6 possible base substitution contexts in our human cancer cells following ENU exposure (**Supplemental Table S8**). There should therefore be greater potential to identify a larger number of resistance mutations regardless of their mutation spectrum. Additionally, we compared the mutation spectrum of ENU to that of another mutagen, namely Gamma irradiation. Two cancer cell lines (MSI versus MSS) were irradiated and single cells expanded to clones prior to exome sequencing. We observed that the mutation spectrum differed quite significantly for the two cell lines, suggesting that unlike ENU mutagenesis there is a greater cell line-specific effect on the pattern of mutations observed (**Supplemental Fig. S10**). This would need to be taken into account if gamma irradiation was used for detection of drug resistance genes.

It is important to note that given it is likely that only one allele is mutated for any specific gene, and less than 5% of the ENU mutations are capable of abrogating protein expression (i.e nonsense, frameshift or essential splice site mutations), the screen is strongly biased towards gain-of-function or dominant point mutations. Loss-of-function resistance genes would therefore be better captured through genome-wide CRISPR inactivation screens or to employ ENU in the setting of haploid cell models (although how relevant these are for aneuploidy cancer cells might be a confounding issue).

A major challenge in the interpretation phenotype-directed screening of ENU mutation models is to identify driver mutations from passenger mutations. This is of particular importance in our experiment given ENU mutagenesis generates an average of almost 500 new mutations per drug resistant clone. We posited that in addition to recurrence, evidence that multiple mutations were enriched within the same network or pathway in a mutually exclusive manner would increase the likelihood of these being driver events. There is a long history of using public resources of such networks to identify enriched genes. We developed a new algorithm SLAP-Enrich to model the likelihood of observing a given number of samples with mutations in the pathway under consideration through a Poisson binomial distribution. It is important to note that identifying resistance mutations in such ENU mutagenesis data is heavily dependent on recurrence of the same mutations across multiple samples and therefore identification of rare resistance mutations will be challenging unless massive numbers of resistance clones are generated and sequenced. This is especially pertinent with respect to the 30 drug resistant CCK-1 or NCH-H508 clones where SLAP-Enrich was unable to detect statistical enrichment of mutations in specific pathways. We suggest at least two plausible explanations – one is that these are rare (and therefore not recurrent) mutations and therefore do not cluster into previously characterized pathways interrogated by SLAP-Enrich; the other is that the observed resistance is the result of mutations outside of the coding exome (for example, in enhancer/promoter or untranslated (UTR) regions) and therefore not amenable to detection using this whole exome capture approach. With respect to the former possibility, we re-ran SLAP-Enrich after removing any variants in the previously identified putative resistance genes (*BRAF*, *KRAS*, *NRAS*, *MAP2K1* etc) to increase the power to detect additional enriched pathways; We found 9 pathways that were enriched when considering a far less stringent significance threshold (FDR < 20%). These were dominated by pathways of neurotransmitter signalling (**Supplemental Table S9**). Given what we know today about EGFR signalling, these are not plausible candidates to confer resistance to Cetuximab. Finally, investigating the possibility of non-coding mutations as resistance drivers would require whole genome sequencing of these hypermutated samples (each harboring approx. 50,000 mutations per genome) and the development of statistical approaches for recurrence detection in non-coding regions. Such algorithms are currently under development as part of the PanCancer Analysis of Whole Genomes (http://pancancer.info/) which is undertaking the analysis of >2000 whole genomes. Thus, in due course as whole genome sequencing costs falls and analytical tools are developed for the non-coding genome, these ENU clones that harbor possible non-coding resistance point mutations could be re-sequenced and re-analysed.

Here we establish a model for the use of genome-wide chemical mutagenesis screens to capture the diversity of clinically relevant drug resistance protein-coding mutations in aneuploid cancer cells. As proof of concept, we employed this screen in the setting of an EGFR therapy and colorectal cancer, a disease in which response to such therapy is invariably followed by the acquisition of resistance. Such resistance mechanisms are heavily dominated by point mutations in the MAP kinase signalling pathway and have been extensively validated in patient cohorts (**Supplemental Table S1**) (Yonesaka et al. 2011; Diaz et al. 2012; Misale et al. 2012; Montagut et al. 2012; Bardelli et al. 2013; Bettegowda 2014). We are able to identify all clinically detected resistance mutations to Cetuximab treatment in colorectal cancer, and in addition potential therapeutic avenues to re-sensitize resistant cells. We propose that ENU mutagenesis should be incorporated alongside newer genome-wide CRISPR gene editing technologies in the systematic interrogation of drug resistance given the prevalence (and potential for therapeutic targeting) of point mutations as mediators of resistance in cancer.

## Methods

### Materials

All cell culture was performed in either RPMI or DMEM/F12 medium (according to the supplier’s recommendations) and supplemented with 5% FBS and penicillin/streptavidin. Cells were maintained at 37°C and 5% CO2 during culture. The identity of all cell lines used in this paper was confirmed using a panel of 95 single nucleotide polymorphisms (SNPs) used previously for cell line authentication (Fluidigm, San Francesco, CA).

### Immunoblotting

Differential phosphorylation of proteins in signalling pathways were analysed by western blot. Cells were plated 24 hours prior to drug treatment and incubated for indicated times and concentrations. Adherent cells were then washed with PBS and collected after indicated incubation time with drug using lysis buffer containing 5% β-mercaptoethanol, 150mM NaCl, 50nM Tris pH 7.5, 2nM EDTA pH 8, 25nM NaF, 1% NP-40, protease inhibitors (Roche), phosphatase inhibitors (Roche). Lysates were then normalised after bicinchoninic acid (BCA) assay using lysis buffer. Protein lysates were resolved using SDS page electrophoresis in pre-cast Invitrogen 4-12% Bis-Tris gels and transferred for 12 hours. Primary antibodies: p44/42 MAPK, Phospho-p44/42 MAPK (Thr202/Tyr204) and Akt were sourced from Cell Signalling and phospho-Akt (pS473) was sourced from Invitrogen. Monoclonal β-tubulin was sourced from Sigma (USA).

### Drug sensitivity assays

Cells were seeded in 96-well plates for 6-day assays and 6-well plates for 20-day clonogenic assays. Cells were incubated in drug free media to allow for adherence for 24 hours before the addition of drug at indicated concentrations. Each cell line was seeded to achieve approx. 70% confluency at the end of the assay. Cetuximab was obtained from the Addenbrookes’ Hospital Pharmacy, Cambridge (UK). Trametinib (GSK1120212) and Dabrafenib were obtained from Selleckchem (USA).

### ENU mutagenesis of cell lines

The CCK-81 and NCI-H508 cell lines were incubated in a concentration range of ENU (0-10 mg/ml) and viability measured after 48 hours. A concentration of 0.1 mg/ml was subsequently selected for resistance models as having modest effect on cell viability while still generating a high rate of mutations (**Supplemental Fig. S11**). Cells were incubated in ENU at the indicated concentration for 24 hours before being washed 3 times with PBS and incubated in media for a further 24 hours. Cells were then selected with 10µg/ml Cetuximab 48 hours post-ENU exposure for 8 weeks. Clones were then picked using Scienceware small cloning cylinders (Wayne, NJ) and either transferred to 96 well plates or expanded into large flasks for drug sensitivity assays. DNA was extracted in 96-well plate format using the Agencourt DNAdvance Genomic DNA Isolation kit.

### Gamma radiation of cell lines

The MSI colon cancer cell line HCT116 and the MSS lung cancer line NCI-H3122 were each irradiated with either 1Gy or 10Gy. The following day single cells were flow sorted and expanded as colonies. DNA was extracted from 9 colonies from each cell line and submitted for sequencing.

### Whole Exome Sequencing

Exome sequencing was carried out using the Agilent SureSelectXT Human All Exon 50Mb bait set. 72 clones were DNA extracted and underwent library construction, flow cell preparation and cluster generation according to the Illumina library preparation protocol. We performed 75-base paired-end Illumina sequencing. Read alignment to the reference human genome (GRCh37) was performed using the Burrows-Wheeler Aligner (BWA) (http://bio-bwa.sourceforge.net/) (Li and Durbin 2010). Unmapped reads were excluded from the analysis. The average coverage across CCK-81-and NCI-508-derived clones was 65X and 62X respectively. The matched parental cell lines were sequenced at greater depth (158X in CCK-81 and 144X in NCI-H508).

### Variant Detection

Single nucleotide substitutions were called using the CaVEMan C (Cancer Variants through Expectations Maximisation) algorithm and insertions/deletions were called using split-read mapping implemented in the Pindel algorithm (https://github.com/cancerit). The CaVEMan algorithm only analyses reads that are properly paired and not marked as duplicates. Variants were identified by comparison to a reference single matched sample consisting of a high sequence coverage contemporary parental cell line control.

### Data Filtering To Remove Pre-existing Subclonal Variants

A number of clones shared mutations which were present in a small percentage of reads in their corresponding contemporary parental cell line sequence. These subclonal mutations could confound subsequent pathway analysis by causing enrichment in a pathway due to mutations which were present before ENU treatment but were not called due to their low representation. To overcome this problem, variants were filtered against the deep sequenced contemporary parental control after mutation calling via Caveman and Pindel. The Samtools mpileup algorithm was used to remove any mutations which were present in 0.5% or more reads in the high coverage parental cell line control (http://samtools.sourceforge.net/). The final set of mutations were used to generate an event matrix for all 72 clones (**Supplemental Table S6**) and used as the input file for SLAP-Enrich analysis described below.

### Deciphering mutational signatures branding exome sequences of clones exposed to ENU

The immediate 5 prime and 3 prime sequence context of base substitutions identified across Cetuximab-resistant clones was extracted using the ENSEMBL Core APIs for human genome build GRCh37 and was used to generate mutational catalogues for the downstream analysis. The mutational catalogue of CCK-81 Cetuximab-resistant clones contained 7,198 substitutions, while the NCI-H508 clones contained a total of 23,862 substitutions. Mutational signatures were deciphered separately across both catalogues of mutations, using a previously developed computational framework (Alexandrov et al. 2013b). Briefly, the algorithm identifies a minimal set of mutational signatures that optimally explains the proportions of mutation types found across a given mutational catalogue (i.e. across all substitutions identified in CCK-81 and NCI-H508 clones; **Supplemental Fig. S4);** and then estimates the contribution of each identified signature to a mutation spectra of each sample included in analysis (i.e. to a mutation spectra of each individual clone; see for **NCI-H508** clones - **Supplemental Fig. S5**).

### Sample Level Analysis of Pathway Enrichments (SLAP-Enrich)

The statistical model implemented in SLAP-Enrich, results from a comparison with similar methods, its unique aspects, other possible analytical options that it allows and results from its application to other case studies (doi: https://doi.org/10.1101/077701). A short description of the statistical model implemented in SLAP-Enrich is also included in the Supplementary Methods, together with specifications of input and parameter settings used for the analyses presented in this manuscript. SLAP-Enrich is implemented as an R package and it is publicly available at https://github.com/saezlab/SLAPenrich. The computational pipeline to reproduce the presented results is implemented in the enclosed BrammeldEtAl_analysis.R script (also available in the Supplementary Methods).

Briefly, as a first step, SLAP-Enrich estimates the probability of observing at least one gene belonging to a given pathway mutated in a given sample, based on the length of the total exon blocks of the genes in that pathway, and the sample mutation burden. Once this probability has been estimated for each individual sample, SLAP-Enrich models the likelihood of observing a given number of samples with mutations in the pathway under consideration through a Poisson binomial distribution. This is the discrete distribution of a sum of Bernoulli trials in which the probability of success is not constant. It is used by SLAP-Enrich to compute the deviance of the number of observed samples with mutations in a given pathway from its expectation through a corresponding p-value assignment.

### Identification of drug resistance genes based on the impact of coding mutations

To identify recurrently mutated driver genes, a *d*_N_/*d*_S_ method that considers the mutation spectrum, the sequence of each gene, the impact of coding substitutions (synonymous, missense, nonsense, splice site) and the variation of the mutation rate across genes were used. Owing to the lack of a neutral reference for the indel rate in coding sequences, a different approach was required (Supplementary Methods for details). Given this approach has been developed for the detection of driver genes in large cancer sample datasets, samples sharing more than 3 mutations (i.e. related subclonal populations) were merged to create a single set of variants composed of the union of all observed variants in the similar samples. In the case of the 72 ENU clones, this resulted in the creation of 19 informative samples that were used for the *d*_N_/*d*_S_ analysis.

To detect genes under significant selective pressure by either point mutations or indels, for each gene the P-values from the *d*_N_/*d*_S_ analysis of substitutions and from the recurrence analysis of indels were combined using Fisher’s method. Multiple testing correction (Benjamini-Hochberg FDR) was performed separately for all genes, stratifying the FDR correction to increase sensitivity (Sun et al. 2006). To achieve a low false discovery rate a conservative q-value cutoff of <0.05 was used for significance and considered significant any gene with qmis_sfdr<0.05 OR qglobal_sfdr<0.05. Please see Supplementary Methods for detailed explanations of these methods

### Site-directed mutagenesis of MAP2K1 expression vectors

In order to validate candidate there four drug resistance mutations from the ENU-based forward genetic screen we sought to create mutated vectors to express within Cetuximab sensitive colorectal cell lines. Wild-type construct for MAP2K1 was ordered from Dharmacon (Lafayette, CO) and taken forward for *in vitro* site-directed mutagenesis reactions using the GENEART® Site-Directed Mutagenesis System from Thermo Fisher Scientific (Waltham, MA). To achieve this, two complementary mutagenic oligonucleotide primers were designed (obtained from Sigma-Aldrich (St Louis, MO) and used to generate gene cDNA expression constructs with desired mutations. Mutations were confirmed using Sanger Sequencing, before being delivered into cells using lentiviral infection.

### Plasma DNA sequencing

DNA extraction was performed with QIAmp DNA Mini kit (Qiagen, Hilden, Germany). Library preparation was done with the Oncomine™ Focus Assay (Thermofisher Scientific, Waltham, MA USA) following the manufacturer’s instructions. After barcoding, libraries were equalized to 100pM. The sequencing template was prepared using the IonPGMSequencing 200 Kit v2 and sequenced in an Ion Select 318 chip using the PGM Sequencing 200 Kit v2 with 500 flows. Hotspot mutations in 35 genes were targeted using the Oncomine Focus Assay (ThermoFisher) (**Supplemental Table S7**). Variant Caller v4.0.r73742 was used for variant calling with the Ion Reporter Software. All filtered variants were also analyzed with the Integrative Genomic Viewer (IGV v2.3) software.

## Data Access

All raw sequence data has been deposited in the European Genome-Phenome Archive (EGA, http://www.ebi.ac.uk/ega/), which is hosted at the EBI, under accession numbers EGAS00001001743, EGAS00001001744 and EGAS00001001745. SLAP-Enrich is implemented as a collection of R scripts and functions and it is publicly available at https://github.com/saezlab/SLAPenrich. The computational pipeline to reproduce the presented results is implemented in the BrammeldEtAl_analysis.R script (also available in the Supplemental Methods).

## Acknowledgements

UM was supported by a Cancer Research UK Clinician Scientist Fellowship. IM is supported by a Cancer Research UK Fellowship. FI was supported by the European Bioinformatics Institute and Wellcome Trust Sanger Institute. J.S.B, S.P. and J.Y. are supported by an ERC Synergy Grant.

Author Contributions: J.S.B performed the majority of experiments, analysed the data and contributed to writing the manuscript. M.P, L.G.A, IM and C.M analysed the data. S.B and M.G designed and ran the combination screen. C.M, A.D and B.B carried out and analysed the plasma sequencing. F.I designed the computational methods, analysed the data and contributed to writing the manuscript. J.S.R and M.R.S supervised the analysis. U.M conceived the project, analysed the data and contributed to writing the manuscript.

**Disclosure Declaration** Competing financial interests - none of the authors hold any competing interests.

